# DRP1/DMNL-1-mediated mitochondrial fission augments *Rickettsia parkeri* replication in macrophages

**DOI:** 10.1101/2025.06.25.661470

**Authors:** Natasha Kelly, Elliott Collins, Basel Abuaita, Juan J. Martinez

## Abstract

Pathogenic Spotted Fever Group (SFG) *Rickettsia* species, including *Rickettsia parkeri* replicate in endothelial cells, monocytes, and macrophages *in vitro* and during infections in murine models of disease. We demonstrated that *R. parkeri* survives and proliferates within phagocytes and avoids intracellular killing within lysosomal compartments. We found that infection of human macrophage-like cells with a related SFG *Rickettsia*, *R. conorii*, resulted in a significant increase in mitochondria-associated proteins, suggesting that mitochondrial functions are involved in *Rickettsia* pathogenesis. Several intracellular bacterial pathogens manipulate host cell mitochondrial networks and stimulate mitochondrial fission mediated by a GTP-binding regulatory protein, DRP1/DMNL1, to promote intracellular replication. Here, we investigated the contribution of DRP1 in the growth of *R. parkeri* in macrophages. Murine immortalized bone marrow derived macrophages (iBMDMs) and primary human monocyte derived macrophages were infected with *R. parkeri* and mitochondrial dynamics (fission and network) were assessed by immunofluorescence microscopy. *R. parkeri* proliferated in macrophages, which coincided with a significant increase in mitochondria content and fission compared to uninfected cells. *R. parkeri* infection led to increases in host cell ATP production primarily due to mitochondrial respiration and bacteria were often found co-localized with mitochondrial fragments. Importantly, *R. parkeri* growth was significantly impacted in DRP1 deficient macrophages. These results suggest that the modulation of mitochondria content and dynamics are essential for replication and survival of pathogenic SFG *Rickettsia* species in macrophages and suggest that the metabolic requirements for obligate intracellular pathogens may differ from other pathogenic Gram-negative intracellular bacteria.

## Introduction

*Rickettsia parkeri* is a Spotted Fever Group (SFG) obligate intracellular Gram-negative human pathogen that is transmitted to humans through tick salivary contents during a blood meal. *R. parkeri* rickettsiosis is found primarily in the southeastern United States and is characterized as a less severe, but important form of rickettsiosis in humans compared to Rocky Mountain spotted fever (RMSF) caused by *Rickettsia rickettsii* (1). In addition, a related SFG *Rickettsi*a species, *R.conorii*, is the agent of Mediterranean spotted fever (MSF), which is endemic to Southern Europe, North Africa and India (2).

Early indications of SFG *Rickettsia* infections in humans are unremarkable and include headache, fever, and malaise. Soon after the tick bite, localized replication of rickettsiae at the inoculation site and ensuing tissue damage may give rise to a necrotic lesion, or eschar. Once established in the host, SFG *Rickettsia* were thought to primarily infect the endothelial lining of the vasculature and cause injury to the vascular endothelium and infiltration of perivascular mononuclear cells leading to vasodilation, an increase in fluid leakage into the interstitial space, and a characteristic dermal rash (3, 4). However, recent data from our group and others has clearly demonstrated that other cell types including monocytes, macrophages, neutrophils, lymphocytes, and hepatocytes are heavily colonized during fatal infections in mammals (5–13). Dissemination of the bacteria in the mammalian host is associated with pulmonary edema, interstitial pneumonia, acute renal failure, liver failure, neurological manifestations, and other multi-organ disorders (14).

Broad-spectrum antibiotics such as doxycycline can be used to clinically manage these infections; however, the window of efficacy for these therapies is narrow necessitating the need for the development of more efficacious anti-rickettsial therapies. Therefore, new therapeutic innovations are urgently needed including host-based approaches to control rickettsial infections.

*R. parkeri* infects recognized target cells such as endothelial cells and can infect circulating monocytes in the peripheral blood and in resident macrophages during infections in murine models of disease (6, 7, 15). PMA-differentiated THP-1 cells have been extensively utilized as model human macrophage cell line for previous studies involving intracellular pathogens such as *Legionella pneumophila* (16), *Coxiella burnetii* (17), *Mycobacterium tuberculosis* (18), and *Chlamydia trachomatis* (19). Using PMA-differentiated THP-1 cells as a model macrophage-like cell, we previously demonstrated that several mammalian pathogenic members of the SFG *Rickettsia* including *R. conorii* (20), *R. rickettsii*, *R. parkeri*, *R. akari* and *R. africae* are capable of surviving and proliferating within these cells (21). Species that are not associated with disease in mammals, including *R. bellii* and *R. montanensis* do not replicate in phagocytic cells and are rapidly killed in intracellular compartments containing lysosomal markers (20, 21). These studies further supported the notion that pathogenic *Rickettsia* are not restricted to infecting the vascular endothelium and that other *Rickettsia*-host cell interactions are likely important in pathogenesis.

To initially elucidate host cell pathways that are stimulated by *Rickettsia* species during infection of macrophages, we performed global transcriptome and proteome analyses of *R. conorii*-infected THP-1 macrophages over different time points of infection. These studies revealed that pathways predicted to be involved in establishing a hospitable intracellular environment for growth and survival are modulated to benefit rickettsial growth and survival. Overall, *R. conorii* infection drives the programming of macrophages towards an environment that permits the establishment of a replicative intracellular niche that is characterized in part by an accumulation of several enzymes of the tricarboxylic acid (TCA) cycle, oxidative phosphorylation (OXPHOS) fatty acid β-oxidation as well as several inner and outer membrane mitochondrial transporters. Increases in these mitochondrial proteins suggested that pathogenic SFG *Rickettsia* species may induce a stimulation of *de novo* mitochondrial biogenesis (mitogenesis), an increase in the oxidative capacity of mitochondria and an increase in the β-oxidation of synthesized host cell fatty acids (22, 23). To further support this hypothesis, we previously demonstrated that pharmacological inhibition of host cell *de novo* fatty acid synthesis and reduced expression of mammalian fatty acid synthase (FASN) greatly inhibited rickettsial growth within macrophages. In addition, pharmacological inhibition studies targeting key mammalian cell enzymes and pathways involved in lipid metabolism including triglyceride lipases (inhibited by Orlistat) and fatty acid β-oxidation (inhibited by Etomoxir) revealed that rickettsial growth was significantly diminished in these cells. Taken together, these studies suggested that the modulation of mitochondrial function to potentially provide ATP through OXPHOS and other important mitochondria-derived metabolites are critical events involved in the growth and survival of pathogenic *Rickettsia* species in mammalian phagocytic cells (22–25).

While the important role(s) of modulating mitochondria function in macrophages by several intracellular pathogens including *C. trachomatis*, *S. typhimurium*, *B. abortus*,

*L. pneumophila* and *L. monocytogenes* has been well established (26), very little in known regarding the impact of these processes to the growth and survival of *Rickettsia* species and other obligate intracellular bacteria in macrophages. Murine immortalized bone marrow derived macrophages (iBMDMs) exhibit characteristics of freshly isolated bone marrow derived macrophages (BMDMs) *in vitro*, can be propagated indefinitely using the appropriate growth media, have been successfully utilized to study pathogen-macrophage interactions for several intracellular bacterial pathogens (27). Using iBMDMs as a macrophage model and primary human monocyte derived macrophages, we have shown that *R. parkeri* can efficiently proliferate within these cells. *R. parkeri* infection in iBMDMs drives an increase in mitochondria content and a significant stimulation of ATP production primarily through mitochondrial respiration. We further demonstrate that infection of iBMDMs and primary human macrophages results in a significant disruption of mitochondrial networks, resembling mitochondrial fission. In addition, *R. parkeri* infection of iBMDMs leads to the hyper serine phosphorylation of a key regulator of mitochondrial dynamics (DRP1/DNM1L) suggesting that activation of mitochondrial fission may be a key event in the infection process. Indeed, inhibition of *drp1* expression in iBMDMs diminishes the ability of *R. parkeri* to efficiently proliferate within these cells. Together these results strongly indicate that DRP1-mediated mitochondrial dynamics play a critical role in the proliferation of *R. parkeri* during infection of mammalian macrophages.

## Materials and Methods

### Cell lines, *Rickettsia* Growth and Purification

African green monkey kidney epithelial cells (Vero) and L929 fibroblast cells were grown at 37^0^C/5%CO_2_ in Dulbecco’s modified Eagle’s medium (DMEM; Corning) supplemented with 10% heat-inactivated fetal bovine serum (hiFBS, Biowest), 1x non-essential amino acids (Corning), and 0.5 mM sodium pyruvate (Corning). Murine immortalized bone marrow derived macrophages (iBMDM) were grown in macrophage media (DMEM supplemented with 20% hiFBS and 30% filtered L929 supernatant) as previously described (27). Lentiviral knockdown of *drp1* expression in iBMDMs and isolation of cell clones has been previously described . *Rickettsia parkeri* isolate Portsmouth was obtained from Dr. Christopher Paddock (US Center for Disease Control) (28), propagated in Vero cells in a humidified 5% CO_2_ incubator at 34^0^C and purified as previously described (29–31). *R. parkeri* was used between passage 2 and passage 4 after being received in our laboratories at the LSU School of Veterinary Medicine.

### Differentiation of human primary macrophages

Human blood samples were obtained from healthy volunteers from the clinical laboratory of Pennington Biomedical Research Center. The mononuclear cells (PBMCs) fraction was isolated by centrifuging the blood onto a Ficoll-Paque plus density gradient medium (Millipore Sigma, Cat# 17-1440-02). PBMCs were incubated at 37°C in 5% CO_2_ and non-adherent cells were removed by extensive washing with PBS. Adherent cells were used to differentiate monocytes into macrophages by culturing the cells in purified human recombinant M-CSF (50 ng/ml, Biolegend, Cat# 574816) in DMEM supplemented with 20% heat-inactivated fetal bovine serum (FBS) for 6 days. Differentiated macrophages were used at 80% confluency, which was estimated to be 10^5^ cells/well in 24 well plates.

### Antibodies

Anti-Rc_PFA_ is a rabbit polyclonal antibody that recognizes multiple species of SFG rickettsiae, including *R. parkeri*, and was previously generated and described (5, 32). Anti-Drp1 rabbit monoclonal antibody (cloneD6C7) was purchased from Cell Signaling Technologies. Anti-Drp1^Ser616^ rabbit polyclonal antibody and anti-Complex I mouse monoclonal antibody (clone 18G12BC2) were purchased from Invitrogen. Alexa Fluor^TM^ 488-conjugated goat anti-rabbit IgG, Alexa Fluor^TM^ 546-conjugated goat anti-rabbit IgG, Alexa Fluor^TM^ 546-conjugated goat anti-mouse IgG, Alexa Fluor^TM^ 647-conjugated goat anti-mouse IgG, Alexa Fluor^TM^ 546-phalloidin, Alexa Fluor^TM^ 488 phalloidin, and DAPI (4’, 6’-diamidino-2-phenylindole) were purchased from Thermo Scientific. Mouse monoclonal anti-β-actin antibody (clone AC-74), anti-mouse IgG-HRP conjugate and anti-rabbit IgG-HRP conjugate were purchased from Sigma Aldrich.

### Analysis of *R. parkeri* Growth Dynamics

Analysis of *R. parkeri* growth in macrophages was performed as previously described with slight modifications (20). Briefly, wild type iBMDMs, *drp1* silenced iBMDMs or primary human macrophages were seeded into 24-well plates in triplicate at a density of 5 x 10^5^ per well. On the day of the experiment, cells were washed once in 1X Dulbecco’s modified phosphate buffered saline (DPBS) and infected with *R. parkeri* at a multiplicity of infection (MOI) of 5 in prewarmed macrophage media. Plates were centrifuged at 300 x g for 5 minutes at room temperature to induce contact between rickettsiae and host cells and then incubated at 34^0^C in the presence of 5% CO_2_ for the indicated time points. Cells were harvested by scraping into 1.0ml of PBS and centrifuged to pellet cells at 900 x g for 5 minutes at room temperature. Total genomic DNA extractions were performed using the PureLink® Genomic DNA Mini Kit (Thermo Fisher Scientific) following the manufacturer’s instructions. Growth of *R. parkeri* species determined via a quantitative PCR (qPCR) assay using a LightCycler 480 II (Roche) utilizing the PerfeCTa FastMixII system (QuantaBio) and the following parameters: 10 minutes at 95^0^C; 50 cycles of 95^0^C for 10 seconds, 58^0^C for 1 minute, and 72^0^C for 1 second. This was followed by a cool-down cycle lasting 30 seconds at 40^0^C. Growth was assessed following the amplification of the rickettsial *sca1* gene and the mammalian *actin* gene by calculating the ratio of *sca1* to *actin* in each well as previously described (20). All experiments were performed using triplicate samples per condition and at least three independent biological experiments.

Immunofluorescence microscopy analysis was also used to verify rickettsial growth within mammalian cells as previously described (20). Briefly, control iBMDMs, *drp1* knockdown iBMDMs or primary human macrophages cells were seeded at a density of 5 x 10^5^ cells on sterilized glass coverslips in 24-well plates . Cells were infected with *R. parkeri* as described above, were performed as described above, washed with 1X PBS and subsequently fixed with 4% PFA in PBS for 20 minutes. *R. parkeri* cells within macrophages were visualized by incubation in antibody incubation buffer (PBS/2% bovine serum albumin (BSA)) containing anti-Rc_PFA_ (1μg/ml), followed by incubation with either Alexa Fluor^TM^ 488-conjugated goat anti-rabbit IgG (1:1000) or Alexa Fluor^TM^ 546-conjugated goat anti-rabbit IgG (1:1000). Nuclei and the actin cytoskeleton were visualized by further incubation with DAPI (1:1000) and Alexa Fluor^TM^ 488-conjugated phalloidin (1:250) or Alexa Fluor^TM^ 546 phalloidin (1:250), respectively. All coverslips were then washed in 1X PBS and then mounted onto glass slides using Mowiol mounting medium. Cells were then viewed and imaged using a Olympus Fluoview FV10i confocal microscope with a 40x oil immersion objective. Images were further processed using Image J software.

### Flow cytometry

Primary human macrophages, wild type iBMDMs and *drp1* knockdown iBMDMs were seeded on 6-well plates at a density of 1 x 10^6^ cells/well and incubated for 24 hours at 37^0^C/5% CO_2._ Triplicate wells were left un-infected or infected with *R. parkeri* at a MOI of 5 for 4, 24, and 48 hrs. After each time point, cells were washed in 1X PBS and each well was scraped into 1.0ml of 1XPBS. Cells were pelleted by centrifugation at 900xg at room temperature and then incubated in 250μl of CytoFix/CytoPerm (BD Biosciences) at 4^0^C for 20 minutes to fix cells. After fixation, cells were washed twice in 1X Perm/Wash buffer (BD Biosciences) and stored in buffer at 4^0^C until processing for flow cytometry analysis. To detect *R. parkeri* in infected cells, control uninfected and *R. parkeri* infected cells were incubated with 1μg/ml of Anti-Rc_PFA_ antibody for 1 hour at room temperature in 1X Perm/Wash buffer. Cells were washed 3 x in 1X Perm/Wash buffer and then incubated for an additional hour at room temperature in 1X Wash/Perm buffer containing Alexa Fluor^TM^ 488-conjugated goat anti-rabbit IgG (1:1000). Cells were washed in 1X Perm/Wash buffer and analyzed on a BD LSR Fortessa X-20 cytometer equipped with BD FACS DIVA analysis software. At least 10,000 cells were analyzed per sample and the experiment was performed at least two independent times.

### Confocal Microscopy analysis of mitochondria in iBMDMs

Primary human macrophages and iBMDMs were seeded at 5 x 10^5^ cells per well in 24-well plates on sterile glass cover slips, allowed to adhere for 24 hours at 37^0^C/5% CO_2_ and then infected with *R. parkeri* at a MOI of 5. Uninfected control cells and *R. parkeri*-infected cells were centrifuged at 300 x g for 5 minutes at room temperature to induce *Rickettsia*-host cell contact, and incubated for 24 hours at 34^0^C/5% CO_2_.

Mammalian cells were washed with 1x PBS and then fixed in 4% paraformaldehyde (PFA) for 20 minutes prior to staining. Cells were permeabilized with PBS/0.05% Triton X-100 for 5 minutes at room temperature and incubated with the primary antibodies rabbit anti-Rc_PFA_ (1μg/ml) and mouse anti-Complex I (1μg/ml) followed by the secondary antibodies Alexa Fluor 546-conjugated goat anti-rabbit IgG (1:1000) and Alexa Fluor 488-conjugated goat anti-mouse IgG (1:1000) and DAPI for iBMDMs and Alexa Fluor 546-conjugated goat anti-rabbit IgG (1:1000), Alexa Fluor 647-conjugated goat anti-mouse IgG (1:1000), Alexa Fluor 488-conjugated phalloidin (1:250) and DAPI (1:1000) for primary human macrophages. The coverslips were mounted onto slides using Mowiol mounting medium, and images were obtained with the use of a Olympus Fluoview FV10i confocal microscope with a 40x or 100x oil immersion objective. Subsequent image processing was done with the ImageJ software.

### Western immunoblotting and DRP1 immunoprecipitation

Wild-type IBMDMs and *drp1* knockdown iBMDMs were seeded in 6 well plates at 1x10^6^ cells/well in macrophage growth media. Cells were washed twice in 1X PBS and 1% NP-40 detergent soluble lysates were generated as previously described. Proteins were quantified by BCA assay (Thermo Scientific), normalized for protein content and then resolved on a 4%-20% mini-protein TGX gradient gel (BioRad). Proteins were transferred to nitrocellulose, blocked in tween-tris buffered saline (1XTBST) containing 2% BSA before incubation with anti-Drp1 rabbit monoclonal antibody (2μg/ml) and anti-rabbit IgG-HRP conjugate (1:25,000). After extensive washing in 1X TBST, membranes were incubated with SuperSignal West Pico chemiluminescence horseradish peroxidase substrate (Thermo Scientific) and then exposed to film. Membranes were stripped with Restore^TM^ western blot stripping buffer (Thermo Scientific), blocked again in 1XTBST/2%BSA and reprobed with mouse anti-β-actin (1μg/ml) and anti-mouse IgG-HRP conjugate (1:25,000) to normalize protein content.

For immunoprecipitation experiments, *R. parkeri* infections were carried out as described above at an MOI of 5 for the indicated time points. Cells were washed in PBS and 1%NP-40 detergent soluble lysates were generated from uninfected and infected cells as previously described (33). Detergent soluble proteins were quantified by BCA assay, normalized for protein content and immunoprecipitated using 2μg/ml rabbit anti-DRP1 (clone D6C7) and 25μl of 50% Protein A/G agarose. Immunoprecipitation reactions were washed in 1% NP-40 buffer and boiled in 25μl of 2X SDS-PAGE sample buffer. Proteins were resolved on 4%-20% mini protein TGX gradient gels (BioRad), transferred to nitrocellulose membranes and then blocked with 1X Tris buffered saline (TBST) 2% BSA before incubation with anti-DRP1^ser616^ (2μg/ml) and anti-rabbit IgG-HRP conjugate (1:25,000). Protein were revealed by incubation of membranes with SuperSignal West Pico chemiluminescence horseradish peroxidase substrate (Thermo Scientific) and exposure to film. Membranes were stripped with Restore^TM^ stripping buffer, blocked again in 1XTBST/2%BSA and reprobed with anti-DRP1 (clone D6C7, 2μg/ml) and anti-rabbit IgG-HRP conjugate (1:25,000) to ensure equal protein content in each immunoprecipitation reaction. Depicted western blots are representative of at least two independent biological experiments.

### Seahorse Analysis

The rate of ATP production from glycolysis (glycoATP) and mitochondrial respiration (mitoATP) were measured by Seahorse XF Real-Time ATP Rate Assay (Agilent, Cat#

103677-100) as previously described (27). NT-control and Drp1 KD macrophages were seeded onto Seahorse XF96 cell culture microplates at a density of 4 X 10^4^ per well and cultured at 37℃ for 24 hrs. Media was replaced and macrophages were left untreated (Mock), stimulated with lipopolysaccharide (LPS; 500 ng/ml) for 6 hrs, infected with Rp for 6 hrs, infected with Rp for 24 hrs, infected with Rp for 48 hrs. During the last hour of the experiment, macrophages were washed with the assay culture media and placed in a non-CO_2_ incubator at 37℃ for 45 min. Media were further exchanged with fresh and warmed assay media. Oxygen consumption and extracellular acidification were monitored using the Seahorse XFe96 Analyzer. Data were analyzed by using the XF Real-Time ATP Rate Assay Report Generator to quantify the rate of ATP.

### Mitochondrial DNA quantification

Macrophages were lifted and resuspended into 200 μl of phosphate saline buffer. Total DNA was isolated using DNeasy Blood and Tissue Kit (Qiagen). Nuclear and mitochondrial DNA in the samples were quantified by real-time PCR using the following premier: 18S forward (5’-TAGAGGGACAAGTGGCGTTC-3’); 18S reverse (5’-CGCTGAGCCAGTCAGTGT-3’); Cytochrome c oxidase 1 (MT-CO1) forward (5’-GCCCCAGATATAGCATTCCC-3’); MT-CO1 reverse (5’-GTTCATCCTGTTCCTGCTCC-3’).

The abundance of mitochondrial DNA (MT-CO1) levels was normalized relative untreated (Mock) macrophages and presented as ratio. The Ct value of MT-CO1 was divided by the Ct value of 18S for each sample. Then, the relative abundance of mitochondrial DNA (MT-CO1) levels was calculated relative to the average ratio of MT-CO1/Nuclear of uninfected samples.

## Results

### *R. parkeri* proliferates within murine immortalized bone marrow derived macrophages (iBMDMs)

We have previously demonstrated that similar to other related SFG *Rickettsia* species, *R. parkeri* strain “Portsmouth” can efficiently invade into and proliferate within PMA-differentiated THP-1 as a model for mammalian macrophages (21). Murine immortalized bone marrow derived macrophages (iBMDMs) are an established model mammalian macrophage and have been previously utilized for the study of well-established intracellular bacterial pathogens (27). We, therefore, sought to determine whether *R. parkeri* could invade into and proliferate within iBMDMs using flow cytometry-based, quantitative PCR (qPCR)-based and fluorescence microscopy-based growth assays in these cells. As shown in Figure 1A and B, infection of iBMDMs with *R. parkeri* leads to a significant increase in the number of macrophages infected over the time course of the experiment and a corresponding increase in the mean fluorescence intensity of infected cells. We have previously established that an increase in fluorescence signal specific to SFG *Rickettsia* species within mammalian cells correlates with an increase in rickettsial load within infected cells (34, 35). We, then, utilized a qPCR-based growth assay and fluorescence microscopy to confirm *R. parkeri* growth within these cells (Figure 1C and D). We also determined that *R. parkeri* efficiently proliferate within primary human macrophages and that growth was not restricted to macrophages of murine origin (Supplementary Figure 1). Taken together, these results confirm that iBMDMs can be utilized as a model for the study of *Rickettsia*-macrophage interactions and that *R. parkeri* can efficiently proliferate within primary human macrophages.

**Figure 1.**
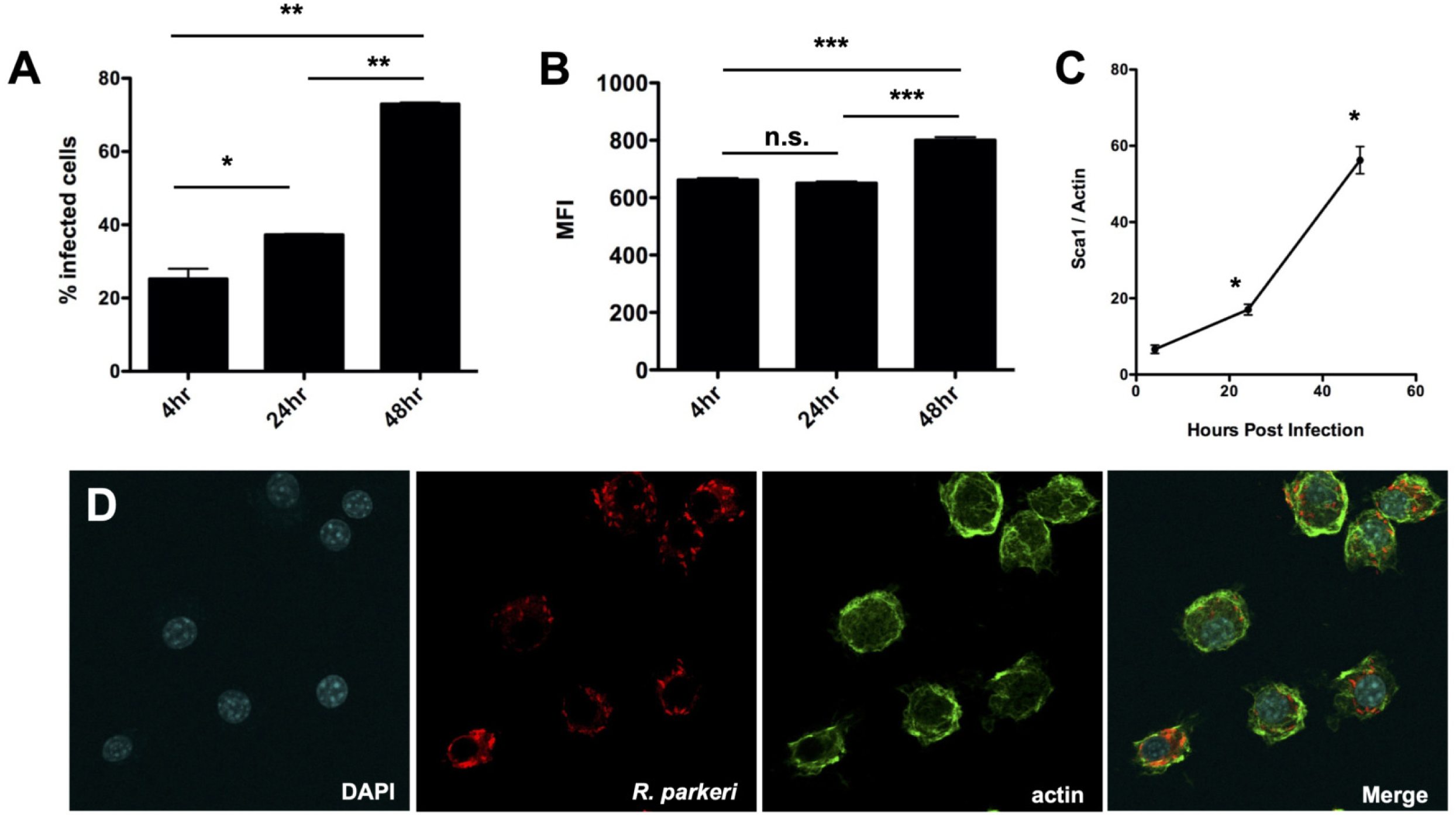
*R. parkeri* strain Portsmouth proliferate in immortalized murine bone marrow derived macrophages (iBMDMs). Flow cytometry analyses demonstrate an increase in the percentage of cells infected and a corresponding increase in the mean fluorescence intensity of *R. parkeri* infected cells over time (A and B). Q-PCR analysis confirms an increase in *R. parkeri* growth as a ratio of rickettsial genomic content (*sca1*) over mammalian cell content (*actin*) (C). *R. parkeri* infected iBMDMs at 48 hours post infection reveal intact bacteria within iBMDMs. Actin (green), *Rickettsia* (red), nuclei (blue). In A and B, *p=0.0132, **p<0.001; in C *p<0.01.

### *R. parkeri* infection of iBMDMs leads to changes in mitochondrial respiration

Previous results from our laboratory demonstrated that infection of differentiated THP-1 macrophage-like cells with a related SFG *Rickettsia* species, *R. conorii*, led to a significant increase in several enzymes involved in the tricarboxylic acid (TCA) cycle, oxidative phosphorylation (OXPHOS), fatty acid β-oxidation and glutaminolysis, as well as of several inner and outer membrane mitochondrial transporters (23). Increases in these mitochondrial proteins suggested that infection of phagocytic cells with a pathogenic SFG *Rickettsia* species would result in a stimulation of *de novo* mitochondrial biosynthesis (mitogenesis) and a corresponding increase in the oxidative capacity of mitochondria in infected cells. To determine if the observed changes in mitochondria-associated protein levels induced by SFG *Rickettsia* infection correspond to an increase in mitochondria content, we infected iBMDMs with *R. parkeri* at an m.o.i=5 for 24 hrs and 72hrs and extracted total DNA from these samples. We quantified mitochondria content in uninfected and infected cells by RT-qPCR analysis of mitochondrial DNA relative to nuclear DNA as described in (36). As shown in Figure 2A, *R. parkeri* infection in iBMDMs leads to a significant increase in mitochondrial DNA abundance relative to uninfected cells at 24 hrs and 72hrs post infection. We further hypothesized that infection of mammalian macrophages by pathogenic *Rickettsia* species will result in significant and dynamic changes in ATP production to potentially fulfill metabolic requirements of this class of obligate intracellular pathogens. We infected iBMDMs at an m.o.i of 5 for 6, 24 and 48 hrs and utilized the Real-Time ATP Rate Assay measured by an Agilent Seahorse XF Pro. As a control, we independently incubated uninfected iBMDMs with LPS (100ng/ml) to stimulate immunometabolic shifts. As shown in Figure 2B, *R. parkeri* induced significant ATP production via glycolysis (glycoATP) similar to the LPS-stimulated control cells and greater than uninfected (mock) control cells. In contrast, ATP production via mitochondrial respiration (mitoATP) was significantly induced by *R. parkeri* infection compared to uninfected cells. LPS-treatment did not induce significant mitoATP levels compared to mock. Mitochondrial-mediated ATP production was stimulated as early as 6 hrs post infection and remained elevated at 24hrs and 48hrs post infection, suggesting that *R. parkeri* infection in macrophages leads to a selective increase in mitochondrial metabolism to establish a niche suitable for replication.

**Figure 2.**
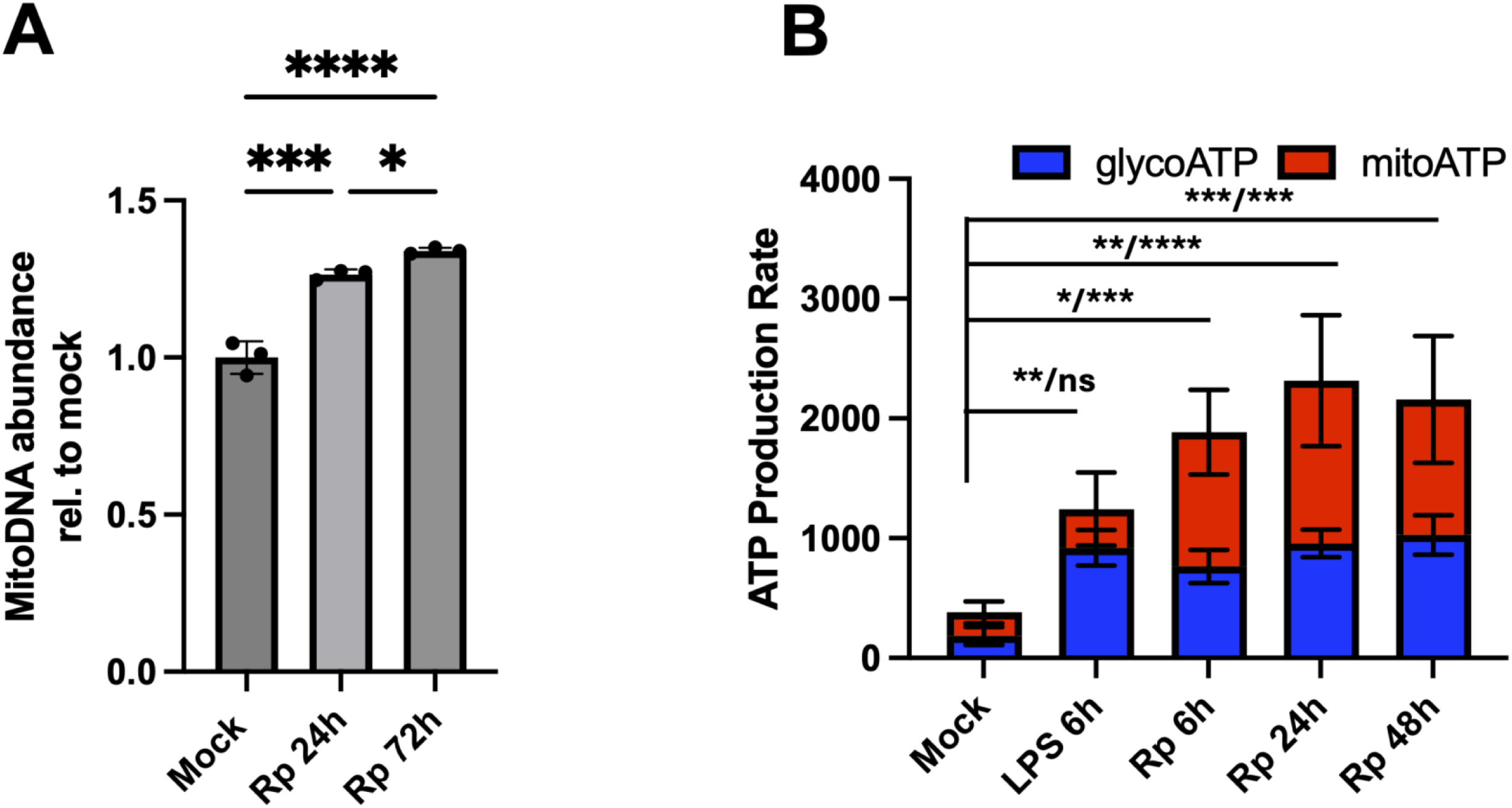
*R. parkeri* infection drives *de novo* mitochondrial biosynthesis and the stimulates increases in ATP production via mitochondrial respiration. A. qPCR analysis of uninfected (mock) or *R. parkeri* infected cells at 24hrs (Rp 24h) and 72hrs (Rp 72h) reveals a significant increase in mitochondrial DNA ANOVA analysis, *p=0.0286, ***p=0.0001 and ****p<0.0001). B. Seahorse analysis of ATP production revealed significant changes in both glycolysis (Blue) and mitochondrial respiration (Red) at indicated times post infection. LPS drives maximal ATP production via glycolysis and is used as a control. *p=0.0345, **p=0.0017, ***p=0.0001, ****p<0.0001, ns= not significant.

### *R. parkeri* infection drives disruption of mitochondrial networks

Several intracellular bacterial pathogens, including *L. pneumophila* (vacuolar) and *L. monocytogenes* (cytoplasmic), manipulate the infected host cell mitochondrial network to promote intracellular replication. The function of mitochondria is influenced by changes in organelle morphology, quantity and often localization. A key regulator of mitochondria morphology is the dynamin 1-like protein (DNM1L or DRP1) which is a GTPase that oligomerizes at the outer mitochondrial membrane participates in the fragmentation of mitochondrial filaments (fission) (37). In the case of *L. pneumophila*, this species uses its type IV secretion machinery to secrete the effector proteins (MitF/LegA), inducing DRP1 dependent mitochondrial fragmentation and ultimately aiding in the formation of a replicative intracellular niche (38). *L. monocytogenes* induces LLO-dependent mitochondrial fragmentation and mitophagy to evade killing within infected mammalian cells (39). To determine if pathogenic *Rickettsia* species can also stimulate changes to mitochondrial networks, we infected murine iBMDMs and primary human macrophages with *R. parkeri* for 24 hours and then processed these samples for analysis by immunofluorescence microscopy. As shown in Figure 3, *R. parkeri* infection of iBMDMs macrophages led to the disruption of the mitochondrial network similar to the mitochondria fragmentation (fission) that had been previously observed for *L. pneumophila* in macrophages (38). In some instances, *R. parkeri* cells co-localize with mitochondrial fragments (Figure 3B arrows). Similar results were observed in primary human macrophages incubated with *R. parkeri* at 24 hrs post-infection (Supplementary Figure 2). These results suggest that modulation of mitochondria content and dynamics may also play a critical role in replication of SFG *Rickettsia* species in mammalian macrophages.

**Figure 3.**
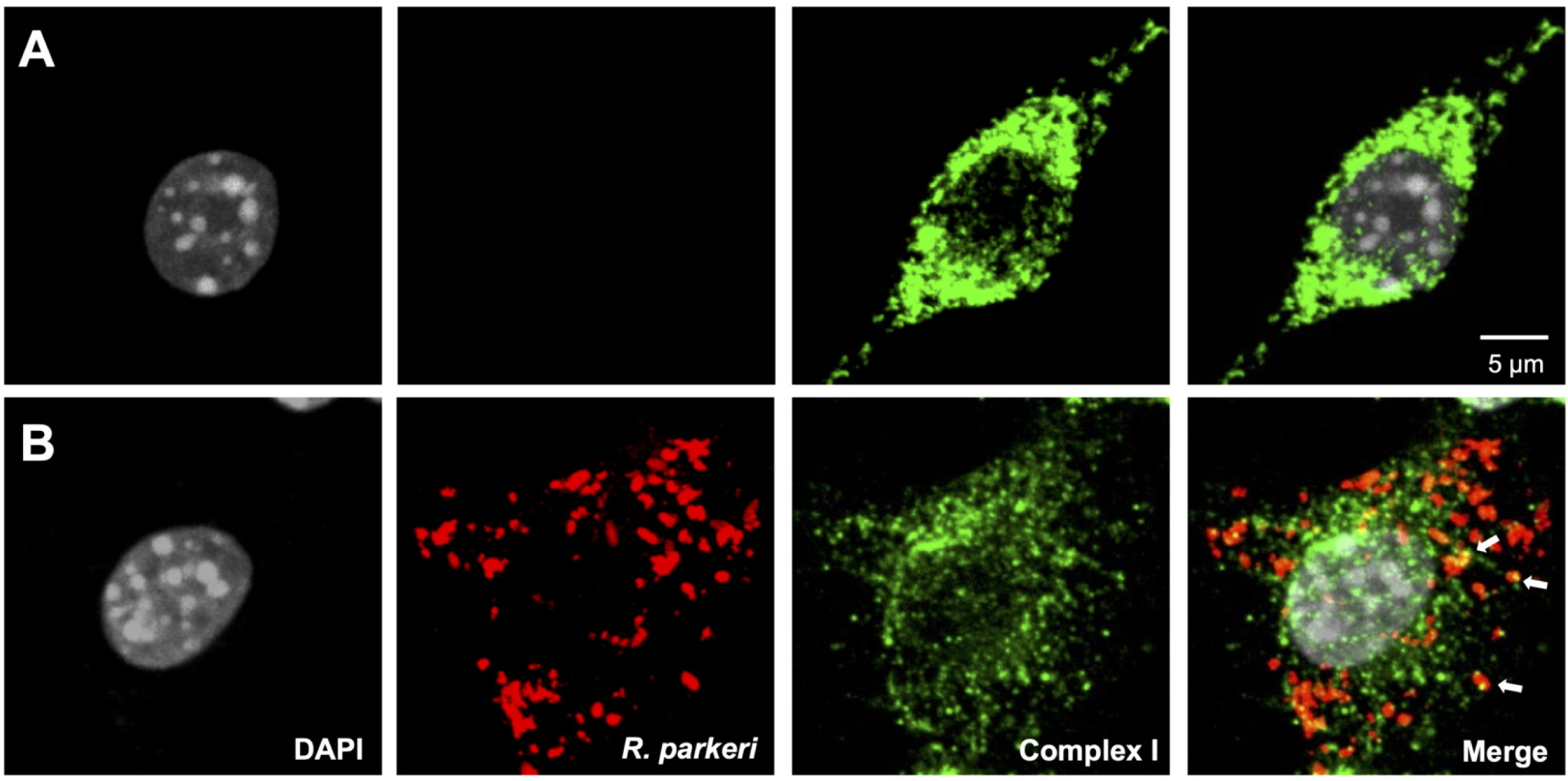
*R. parkeri* induced mitochondrial dynamics in mammalian macrophages. Infection of iBMDMs drives significant changes in mitochondria morphology at 24 hrs post infection. A) uninfected (mock, top panels) and *R. parkeri* infected cells (bottom panels). Cells were processed for immunofluorescence microscopy using anti-*Rickettsia* antibody (red), anti-complex I mAb (green) and DAPI, (white). Arrows in B (merge) indicate areas of colocalization between intact *R. parkeri* cells and fragmented mitochondria.

### *R. parkeri* infection of iBMDMs results in DRP1 serine phosphorylation

Activity of DRP1/DNM1L is stimulated by phosphorylation of serine residues including Ser616, which can be assayed by western immunoblotting (38). To determine whether *R. parkeri* infection can stimulate DRP1 activation, we infected iBMDMs for the indicated time points, solubilized mammalian proteins and analyzed the DRP1 phosphorylation state using immunoprecipitation and western immunoblotting with commercially available DRP1-antibodies and DRP1-phosphospecific antibodies. As shown in Figure 4, infection of iBMDMs leads to a rapid increase in DRP1-Ser616 phosphorylation that is maintained up to 48 hours post-infection. These results demonstrate that *R. parkeri* can stimulate DRP1 activity during infections of macrophages.

**Figure 4.**
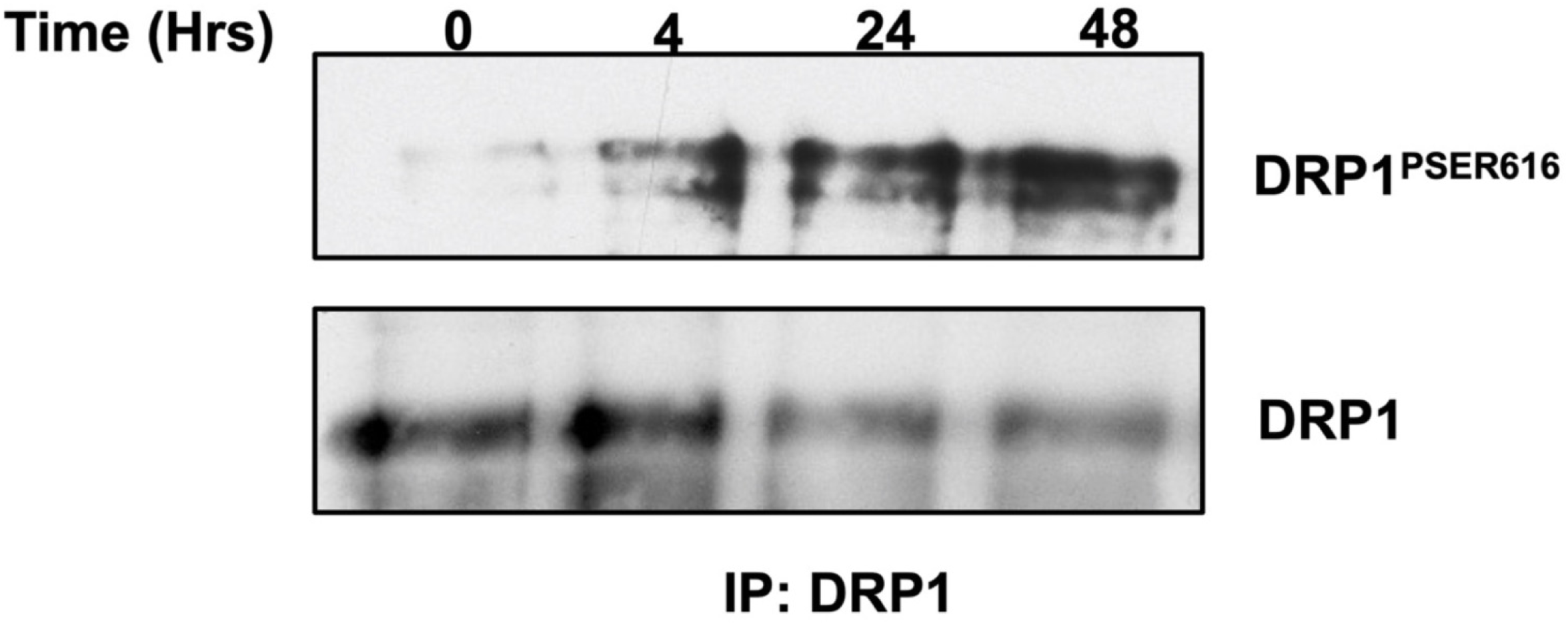
*R. parkeri* stimulates DRP1 serine phosphorylation in iBMDMs. Macrophages were infected for the indicated time points with *R. parkeri* at an moi=5. Detergent soluble protein lysates were generated and equal amounts of total proteins were immunoprecipitated with an anti-DRP1 antibody. Transferred proteins were probed with an anti-DRP1^pSer616^ antibody and re-probed with an anti-DRP1 antibody to control for protein loading.

### Inhibition of DRP1 expression results in decreased *R. parkeri* growth in macrophages

Previous studies have demonstrated that infection of human monocyte derived macrophages by wild type *L. pneumophila* led to a significant increase in the serine phosphorylation of DRP1. In addition, pharmacological inhibition of DRP1 in primary macrophages or in cells silenced for DRP1 expression, reduced intracellular replication of *L. pneumophila* (38) and *L. monocytogenes* (39), suggesting that mitochondrial fragmentation/fission can favor bacterial replication. We hypothesized that the observed dynamic changes to mitochondria and DRP1 Ser616 phosphorylation in *R. parkeri*-infected macrophages would also favor the establishment of a replicative intracellular niche. To determine whether DRP1 activity plays a role in *R. parkeri* replication in macrophages, we silenced *drp1* expression in iBMDMs by shRNA lentiviral transfection as previously described (27). Transfection of iBMDMs with *drp1* specific shRNAs effectively reduced DRP1 expression compared to control transfected cells as determined by western immunoblotting (Figure 5A). We next infected control and *drp1*-knockdown iBMDMs with *R. parkeri* at an moi of 5 for 4, 24 and 48hrs and analyzed proliferation by flow cytometry, q-PCR and fluorescence microscopy. As shown in Figure 5B-E, *R. parkeri* growth was significantly reduced in *drp1* knockdown cells compared to control transfected cells. Taken together, these results demonstrate that DRP1 activity and expression are required for efficient *R. parkeri* proliferation within infected mammalian macrophages.

**Figure 5.**
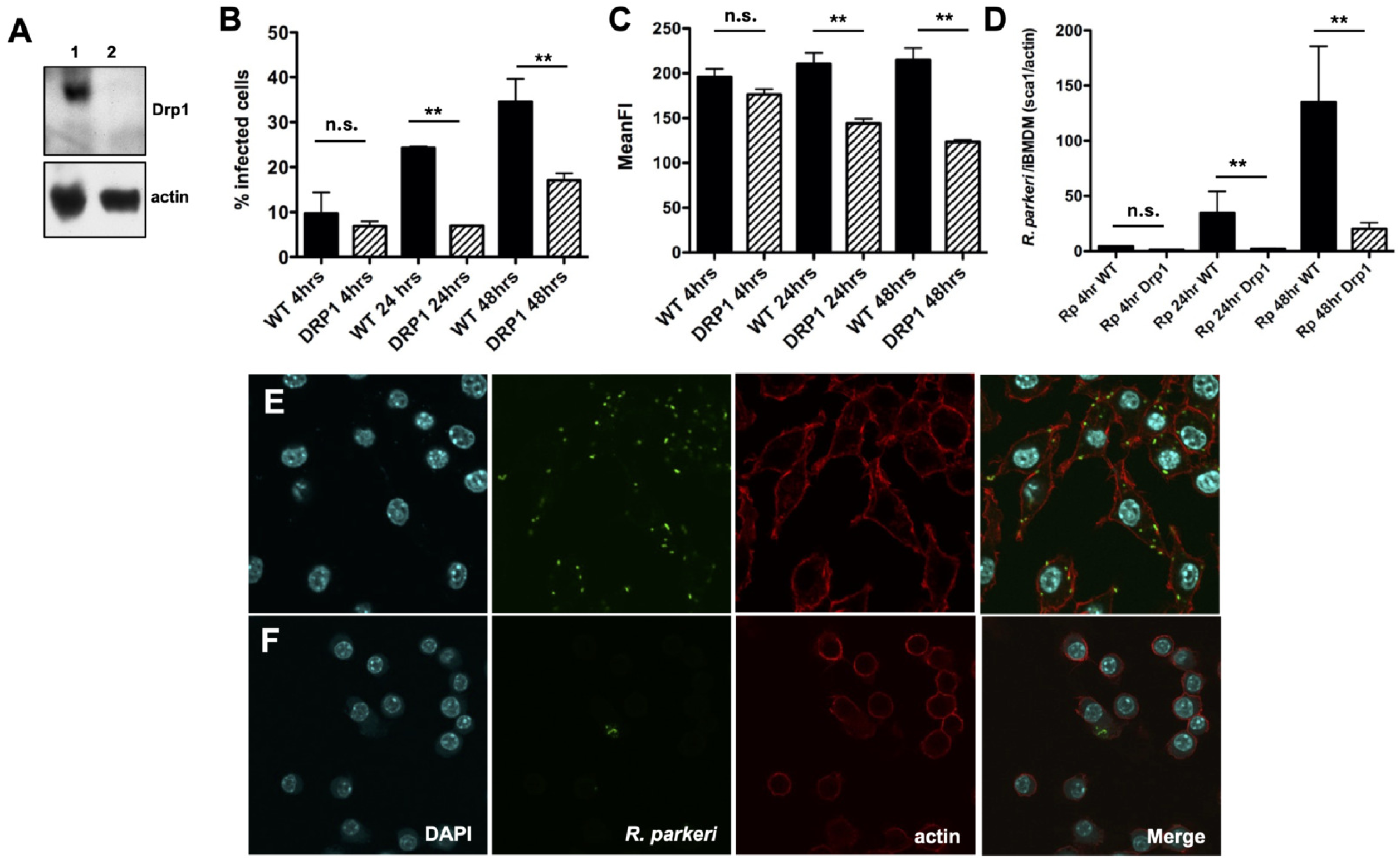
Inhibition of DRP1 expression in iBMDMs diminishes *R. parkeri* proliferation. A) Western immunoblot ananlysis of generated detergent soluble lysates from control (1) and *drp1* knockdown cells (2) reveals significant inhibition of DRP1 protein expression. B and C) Flow cytometry analysis of wild type (WT) and *drp1* knockdown (DRP1) infected cells and the mean fluorescence intensity (MFI) of *R. parkeri* indicated time points. D quantitative PCR (qPCR) analysis of WT and DRP1 iBMDMs infected with *R. parkeri*. E and F) Immunofluorescence microscopy analysis of *R. parkeri* WT-infected iBMDMs (E) and DRP1-infected iBMDMs (F) demonstrates a significant growth inhibition in DRP1 knockdown cells.

## Discussion

As obligate intracellular pathogens, several *Rickettsia* species have evolved strategies to infect different mammalian cell types including endothelial cells, monocytes and macrophages and to sequester nutrients within these infected cells required to establish productive replicative niches. Comparative genomic analyses between pathogenic and non-pathogenic *Rickettsia* species have thus far not revealed “smoking guns” which could explain why one *Rickettsia* species can cause disease in a mammal, while another is essential avirulent. (40, 41). Our recent studies have demonstrated that the ability of a *Rickettsia* species to cause disease in mammals is often correlated with the ability of that species to infect and proliferate within phagocytic cells. For example, we demonstrated that recognized pathogenic species including *R. parkeri*, *R. rickettsii*, *R. conorii*, *R. akari* and *R. africae* can effectively avoid lysosomal compartment within PMA-differentiated THP-1 cells (macrophages) and efficiently proliferate within these cells while two species that are not associated with disease in mammals, *R. bellii* and *R. montanensis*, are rapidly killed in compartments containing lysosomal markers. However, the molecular mechanisms that promote rickettsial survival and growth within mammalian phagocytic cells have yet to be fully elucidated.

Our previous work established that *R. conorii* and likely other related *Rickettsia* pathogenic species reprogram signaling networks and pathways within infected macrophages to aid in the establishment of a permissive replicative intracellular niche. (22, 23). Studies from our group clearly demonstrated that infection of THP-1 macrophages with *R. conorii* led to the predicted upregulation of pathways associated with *de novo* host cell fatty acid biosynthesis and b-oxidation of these fatty acids, presumably for the generation of ATP and changes in mitochondria abundance and function. Indeed, infection of THP-1 macrophages with *R. conorri* led to a significant increase in the abundance of mitochondrial transporters including voltage dependent ion channels 1-3 (VDAC1-3), solute carrier family 25 members 1, 3, 5, and 6 (SLC25A1, 3, 5, 6), translocase of outer mitochondrial membrane 22 and 40 (TOMM 22 and TOMM44) along with an increase in proteins from Complex I (NDUFB8), Complex II (SDHB), Complex III (UQCRC2) and Complex IV (CoxII). In addition, rickettsial infections of THP-1 macrophages also led to a decrease in the abundance of several enzymes involved in host cell glycolysis, including phosphoglycerate kinase 1 (PGK1), pyruvate kinase (PKM) and enolase 1 (ENO1), strongly suggesting that rickettsial infections reprogram host cell metabolism to favor oxidative phosphorylation (OXPHOS) over glycolysis. Furthermore, pharmacological inhibition of host fatty acid synthase (FASN), host triglyceride lipases and host fatty acid β-oxidation significantly diminishes intracellular growth (20, 22), highlighting the importance of these processes to rickettsial infections of host target cells. In this current study, we demonstrated that infection of mammalian macrophages (iBMDMs) with *R. parkeri* also leads to a significant increase in the quantity of mitochondria during the time course of the experiment and this induced change in mitochondrial content also coincided with an increased synthesis of host cell ATP primarily through mitochondrial respiration. In addition, *R. parkeri* induced stimulation of mitogenesis and a commensurate increase in the oxidative capacity of mitochondria in infected cells strongly implicates that stimulating *de novo* synthesis of host derived fatty acids and targeting of these fatty acids for OXPHOS are critical for efficient *Rickettsia* proliferation within infected mammalian macrophages (22, 23).

Several intracellular pathogens are found in a pathogen-containing vacuole (PCV) that requires host lipids for structural maintenance of their intracellular niche and nutrients that can be safely and efficiently sequestered in the PCV. In contrast, pathogenic *Rickettsia* species are cytosolic intracellular bacteria that escape from transient intracellular membrane-bound compartments and therefore, lack the accessibility of nutrients within a PCV (24). Previous comparative genomic studies of different *Rickettsia* species demonstrated that this class of obligate intracellular bacteria lack a full arsenal of genes encoding enzymes that are vital to synthesize certain fatty acids and to efficiently produce ATP (42, 43), strongly suggesting that *Rickettsia* species must exploit host cell pathways to obtain these and other host-derived nutrients to fulfill metabolic requirements. Interestingly, *L. pneumophila* infection of human monocyte derived macrophages results in an overall increase in host cell glycolysis and a decrease in OXPHOS along with changes in mitochondrial morphology (Warburg-like metabolism) to establish a more replicative favorable intracellular niche (38). Our results strongly suggest that the metabolic requirements between intracellular Gram-negative bacterial pathogens are distinct within macrophages and may be dictated by their respective intracellular localizations.

In this study, we have demonstrated that infection of mammalian macrophages by *R. parkeri* leads to a drastic reorganization of mitochondrial networks in a process resembling mitochondrial fission. Mitochondrial dynamics including mitochondrial fission are governed by DRP1/DNM1L, a small GTP-binding protein that is activated by serine phosphorylation of a key residue (Ser616) (44). We have also demonstrated that *R. parkeri* infection of macrophages resulted in a time dependent induction of Drp1 serine phosphorylation starting at 4hrs post-infection strongly suggesting that Drp1 is activated during the infection. Interestingly, the facultative intracellular Gram-negative pathogen, *L. pneumophila* uses its type IV secretion machinery to secrete the effector protein (MitF/LegA) which is involved in DRP1 dependent mitochondrial fragmentation and ultimately aids in the formation of a replicative intracellular niche (38). Interestingly, several *R. parkeri* secreted proteins termed “secreted *Rickettsia* factors” (SrfsA-G) have recently been shown to interact with several intracellular compartments and organelles, including the host cytoplasm, mitochondria and endoplasmic reticulum *in vitro*. One of these proteins, SrfB co-localizes with mitochondrial apoptosis inducing factor (AIF); however, SrfB was not sufficient to mediate significant changes to the mitochondrial networks (45) as observed in our study. In addition, genomic and predicted proteomic bioinformatics analyses have yet to reveal a *mitF*/*legA* gene or protein homologue annotated within *Rickettsia* genomes suggesting that either induced mitochondrial fragmentation is not dependent on this type IV secretion system effector protein or that a *Rickettsia* MitF/LegA homologue has yet to be properly annotated. The rickettsial antigen(s) that are involved in mediating changes to mitochondrial morphology in infected macrophages are not known and are currently under investigation.

We determined that in some cases, *R. parkeri* appeared to co-localize with induced mitochondrial fragments within infected murine and human macrophages. The consequences of these interactions are unclear, but these observations lead to the intriguing possibility that establishing these contacts may be a mechanism by which rickettsiae can access and sequester necessary nutrients and metabolites from the mammalian cell. This idea is not without precedent as other Gram-negative intracellular bacteria, such as *Chlamydia trachomatis* and *Coxiella burnetii* secrete effector proteins that make direct contacts between the pathogen containing vacuole (PCV) in which they reside and target host organelles such as the endoplasmic reticulum in areas of contact termed inter-organelle membrane contact sites (MCS). These contacts are hypothesized to be in part required to acquire nutrients and metabolites into the PCV for growth and inhibition of these contacts negatively impairs intracellular bacterial replication (46, 47). A recent study demonstrated that *R. parkeri* in infected mammalian epithelial and endothelial cells interacts with the ER in MCS governed by VAMP-associated proteins (VAPs). However, inhibition of these MCS induced by *R. parkeri* does not appear to positively or negatively impact rickettsial survival or proliferation within these infected cells (48). Whether establishment of MCS between *Rickettsia* species and mitochondria is required for efficient growth within macrophages and other target mammalian cells is an area of active investigation.

The host cell pathways that are stimulated by *R. parkeri* infection and lead directly to DRP1 activation and mitochondrial fission are not yet elucidated. A recent study implicated key enzymes in the mevalonate pathway as playing key roles in directly influencing mitochondrial dynamics. Nitrogen containing bisphosphonates (NBPs such as alendronate and zolendronate) inhibit farnesyl diphosphate synthase (FPP) and results in the inhibition of the biosynthesis of cholesterol, isoprenoids, heme and ubiquinones (49). Interestingly, targeting of FPP by NBPs led to a decrease in mitochondrial fission markers in mammalian endothelial cells (50). We and others have demonstrated that inhibition of HMG-CoA reductase by statin drugs reduced *R. conorii* (34) and *R. parkeri* (51) growth with infected mammalian cells. These results lead to the intriguing possibility that rickettsial species stimulate upstream signaling pathways leading to DRP1 activation and the observed changes to mitochondrial content and morphology. These results led to the intriguing notion that reducing the capacity to induce mitochondrial fission by targeting HMG-CoA reductase and FPP in infected cells will coincide with a reduced capacity for *Rickettsia* species to proliferate intracellularly. Therefore, targeting pathways leading to mitochondrial fission during infection may represent a novel strategy for therapeutic intervention against pathogenic *Rickettsia* species.

## Acknowledgments

We would like to thank current and former members of the Martinez lab for helpful discussions regarding this work. This work was supported in part by an LSU School of Veterinary Medicine (LSU-SVM) graduate student fellowship and a LSU Pinkie Gordon Lane Graduate School Future Scholars Initiative fellowship to NK. This work was also supported in part by a previous award by the National Institutes of Health, National Institute of Allergy and Infectious Diseases (grant AI072606) to JJM.

---

**Supplemental Figure 1. *R. parkeri* proliferates within primary human monocyte derived macrophages.** A-C) Flow cytometry analysis of *R. parkeri* growth in human primary macrophage demonstrates an increase in *Rickettsia* specific fluorescence (A), the percent of infected cells in the population (B) and the quantification of fluorescent signal within infected cells (C). D) Increases in fluorescence signals are correlated with growth as demonstrated by fluorescence microscopy analysis of *R. parkeri* infected human primary macrophages at 48 hours post-infection. DAPI (blue), *R. parkeri* (red), actin (green). Scale bar =5μm. **p<0.001.

**Supplemental Figure 2. *R. parkeri* stimulates mitochondrial fission within primary human monocyte derived macrophages.** (A) uninfected macrophages display a network of intact mitochondria surrounding the nucleus. (B) Infection by *R. parkeri* leads to disruption of mitochondrial networks resembling mitochondrial fission. *R. parkeri* cells are found co-localized with mitochondrial fragments throughout the cell (arrow). DAPI (blue), *R. parkeri* (red), Complex I (white), actin (green). Scale bar =5μm.

## References

1. Paddock CD, Finley RW, Wright CS, Robinson HN, Schrodt BJ, Lane CC, Ekenna O, Blass MA, Tamminga CL, Ohl CA, McLellan SL, Goddard J, Holman RC, Openshaw JJ, Sumner JW, Zaki SR, Eremeeva ME. 2008. *Rickettsia parkeri* rickettsiosis and its clinical distinction from Rocky Mountain spotted fever. Clinical Infectious Diseases 47:1188–96.

2. de Sousa R, Nobrega SD, Bacellar F, Torgal J. 2003. Mediterranean spotted fever in Portugal: risk factors for fatal outcome in 105 hospitalized patients. Annals of the New York Academy of Sciences 990:285–94.

3. Walker DH, Occhino C, Tringali GR, Di Rosa S, Mansueto S. 1988. Pathogenesis of rickettsial eschars: the tache noire of boutonneuse fever. Human Pathology 19:1449–54.

4. Hand WL, Miller JB, Reinarz JA, Sanford JP. 1970. Rocky Mountain spotted fever. A vascular disease. Archives of Internal Medicine 125:879–82.

5. Chan YGY, Riley SP, Chen E, Martinez JJ. 2011. The molecular basis of immunity to rickettsial infection conferred through outer membrane protein B. Infection and Immunity 79:2303–2313.

6. Riley SP, Cardwell MM, Chan YG, Pruneau L, Del Piero F, Martinez JJ. 2015. Failure of a heterologous recombinant Sca5/OmpB protein-based vaccine to elicit effective protective immunity against *Rickettsia rickettsii* infections in C3H/HeN mice. Pathog Dis 73:ftv101.

7. Riley SP, Fish AI, Del Piero F, Martinez JJ. 2018. Immunity against the Obligate Intracellular Bacterial Pathogen *Rickettsia australis* Requires a Functional Complement System. Infect Immun 86.

8. Riley SP, Fish AI, Garza DA, Banajee KH, Harris EK, del Piero F, Martinez JJ. 2016. Nonselective Persistence of a Rickettsia conorii Extrachromosomal Plasmid during Mammalian Infection. Infect Immun 84:790–7.

9. Bechelli J, Vergara L, Smalley C, Buzhdygan TP, Bender S, Zhang W, Liu Y, Popov VL, Wang J, Garg N, Hwang S, Walker DH, Fang R. 2019. Atg5 Supports *Rickettsia australis* Infection in Macrophages In Vitro and In Vivo. Infect Immun 87.

10. Manor E. 1992. The effect of monocyte-derived macrophages on the growth of *Rickettsia conorii* in permissive cells. Acta Virol 36:13–8.

11. Engstrom P, Burke TP, Mitchell G, Ingabire N, Mark KG, Golovkine G, Iavarone AT, Rape M, Cox JS, Welch MD. 2019. Evasion of autophagy mediated by *Rickettsia* surface protein OmpB is critical for virulence. Nat Microbiol 4:2538–2551.

12. Radulovic S, Price PW, Beier MS, Gaywee J, Macaluso JA, Azad A. 2002. *Rickettsia*-macrophage interactions: host cell responses to *Rickettsia akari* and *Rickettsia typhi*. Infect Immun 70:2576–82.

13. Manor E, Sarov I. 1990. Tumor necrosis factor alpha and prostaglandin E2 production by human monocyte-derived macrophages infected with spotted fever group rickettsiae. Ann N Y Acad Sci 590:157–67.

14. Raoult D, Roux V. 1997. Rickettsioses as paradigms of new or emerging infectious diseases. Clinical Microbiology Reviews 10:694–719.

15. Londono AF, Mendell NL, Walker DH, Bouyer DH. 2019. A biosafety level-2 dose-dependent lethal mouse model of spotted fever rickettsiosis: *Rickettsia parkeri* Atlantic Rainforest strain. PLoS Negl Trop Dis 13:e0007054.

16. Takemura H, Yamamoto H, Kunishima H, Ikejima H, Hara T, Kanemitsu K, Terakubo S, Shoji Y, Kaku M, Shimada J. 2000. Evaluation of a human monocytic cell line THP-1 model for assay of the intracellular activities of antimicrobial agents against *Legionella pneumophila*. J Antimicrob Chemother 46:589–94.

17. Ghigo E, Capo C, Tung CH, Raoult D, Gorvel JP, Mege JL. 2002. *Coxiella burnetii* survival in THP-1 monocytes involves the impairment of phagosome maturation: IFN-gamma mediates its restoration and bacterial killing. J Immunol 169:4488–95.

18. Theus SA, Cave MD, Eisenach KD. 2004. Activated THP-1 cells: an attractive model for the assessment of intracellular growth rates of *Mycobacterium tuberculosis* isolates. Infect Immun 72:1169–73.

19. Carratelli CR, Rizzo A, Catania MR, Galle F, Losi E, Hasty DL, Rossano F. 2002. *Chlamydia pneumoniae* infections prevent the programmed cell death on THP-1 cell line. FEMS Microbiol Lett 215:69–74.

20. Curto P, Simoes I, Riley SP, Martinez JJ. 2016. Differences in Intracellular Fate of Two Spotted Fever Group *Rickettsia* in Macrophage-Like Cells. Front Cell Infect Microbiol 6:80.

21. Kristof MN, Allen PE, Yutzy LD, Thibodaux B, Paddock CD, Martinez JJ. 2021. Significant Growth by *Rickettsia* Species within Human Macrophage-Like Cells Is a Phenotype Correlated with the Ability to Cause Disease in Mammals. Pathogens 10.

22. Curto P, Riley SP, Simoes I, Martinez JJ. 2019. Macrophages Infected by a Pathogen and a Non-pathogen Spotted Fever Group *Rickettsia* Reveal Differential Reprogramming Signatures Early in Infection. Front Cell Infect Microbiol 9:97.

23. Curto P, Santa C, Allen P, Manadas B, Simoes I, Martinez JJ. 2019. A Pathogen and a Non-pathogen Spotted Fever Group *Rickettsia* Trigger Differential Proteome Signatures in Macrophages. Front Cell Infect Microbiol 9:43.

24. Allen PE, Martinez JJ. 2020. Modulation of Host Lipid Pathways by Pathogenic Intracellular Bacteria. Pathogens 9.

25. Allen PE, Noland RC, Martinez JJ. 2021. *Rickettsia conorii* survival in THP-1 macrophages involves host lipid droplet alterations and active rickettsial protein production. Cell Microbiol doi:10.1111/cmi.13390:e13390.

26. Ramond E, Jamet A, Coureuil M, Charbit A. 2019. Pivotal Role of Mitochondria in Macrophage Response to Bacterial Pathogens. Front Immunol 10:2461.

27. Gao F, Reynolds MB, Passalacqua KD, Sexton JZ, Abuaita BH, O’Riordan MXD. 2020. The Mitochondrial Fission Regulator DRP1 Controls Post-Transcriptional Regulation of TNF-alpha. Front Cell Infect Microbiol 10:593805.

28. Paddock CD, Sumner JW, Comer JA, Zaki SR, Goldsmith CS, Goddard J, McLellan SL, Tamminga CL, Ohl CA. 2004. *Rickettsia parkeri*: a newly recognized cause of spotted fever rickettsiosis in the United States. Clin Infect Dis 38:805–11.

29. Ammerman NC, Beier-Sexton M, Azad AF. 2008. Laboratory maintenance of *Rickettsia rickettsii*. Current Protocols in Microbiology Chapter 3:Unit 3A 5.

30. Chan YG, Cardwell MM, Hermanas TM, Uchiyama T, Martinez JJ. 2009. Rickettsial outer-membrane protein B (rOmpB) mediates bacterial invasion through Ku70 in an actin, c-Cbl, clathrin and caveolin 2-dependent manner. Cell Microbiol 11:629–44.

31. Chan YG, Riley SP, Chen E, Martinez JJ. 2011. Molecular basis of immunity to rickettsial infection conferred through outer membrane protein B. Infect Immun 79:2303–13.

32. Cardwell MM, Martinez JJ. 2012. Identification and characterization of the mammalian association and actin-nucleating domains in the Rickettsia conorii autotransporter protein, Sca2. Cell Microbiol 14:1485–95.

33. Martinez JJ, Seveau S, Veiga E, Matsuyama S, Cossart P. 2005. Ku70, a component of DNA-dependent protein kinase, is a mammalian receptor for *Rickettsia conorii*. Cell 123:1013–1023.

34. Czyz DM, Potluri LP, Jain-Gupta N, Riley SP, Martinez JJ, Steck TL, Crosson S, Shuman HA, Gabay JE. 2014. Host-directed antimicrobial drugs with broad-spectrum efficacy against intracellular bacterial pathogens. MBio 5:e01534–14.

35. Riley SP, Macaluso KR, Martinez JJ. 2015. Electrotransformation and Clonal Isolation of *Rickettsia* Species. Curr Protoc Microbiol 39:3A 6 1-3A 6 20.

36. Bronner DN, O’Riordan MX. 2016. Measurement of Mitochondrial DNA Release in Response to ER Stress. Bio Protoc 6.

37. Friedman JR, Lackner LL, West M, DiBenedetto JR, Nunnari J, Voeltz GK. 2011. ER tubules mark sites of mitochondrial division. Science 334:358–62.

38. Escoll P, Song OR, Viana F, Steiner B, Lagache T, Olivo-Marin JC, Impens F, Brodin P, Hilbi H, Buchrieser C. 2017. *Legionella pneumophila* Modulates Mitochondrial Dynamics to Trigger Metabolic Repurposing of Infected Macrophages. Cell Host Microbe 22:302–316 e7.

39. Zhang Y, Yao Y, Qiu X, Wang G, Hu Z, Chen S, Wu Z, Yuan N, Gao H, Wang J, Song H, Girardin SE, Qian Y. 2019. *Listeria* hijacks host mitophagy through a novel mitophagy receptor to evade killing. Nat Immunol 20:433–446.

40. Clark TR, Noriea NF, Bublitz DC, Ellison DW, Martens C, Lutter EI, Hackstadt T. 2015. Comparative genome sequencing of *Rickettsia rickettsii* strains that differ in virulence. Infect Immun 83:1568–76.

41. Ellison DW, Clark TR, Sturdevant DE, Virtaneva K, Porcella SF, Hackstadt T. 2008. Genomic comparison of virulent *Rickettsia rickettsii* Sheila Smith and avirulent *Rickettsia rickettsii* Iowa. Infection & Immunity 76:542–50.

42. Merhej V, Raoult D. 2011. Rickettsial evolution in the light of comparative genomics. Biol Rev Camb Philos Soc 86:379–405.

43. Driscoll TP, Verhoeve VI, Guillotte ML, Lehman SS, Rennoll SA, Beier-Sexton M, Rahman MS, Azad AF, Gillespie JJ. 2017. Wholly *Rickettsia*! Reconstructed Metabolic Profile of the Quintessential Bacterial Parasite of Eukaryotic Cells. MBio 8.

44. Taguchi N, Ishihara N, Jofuku A, Oka T, Mihara K. 2007. Mitotic phosphorylation of dynamin-related GTPase Drp1 participates in mitochondrial fission. J Biol Chem 282:11521–9.

45. Sanderlin AG, Kurka Margolis H, Meyer AF, Lamason RL. 2024. Cell-selective proteomics reveal novel effectors secreted by an obligate intracellular bacterial pathogen. Nat Commun 15:6073.

46. Angara RK, Sadi A, Gilk SD. 2024. A novel bacterial effector protein mediates ER-LD membrane contacts to regulate host lipid droplets. EMBO Rep 25:5331–5351.

47. Cortina ME, Derre I. 2023. Homologues of the Chlamydia trachomatis and *Chlamydia muridarum* Inclusion Membrane Protein IncS Are Interchangeable for Early Development but Not for Inclusion Stability in the Late Developmental Cycle. mSphere 8:e0000323.

48. Acevedo-Sanchez Y, Woida PJ, Anderson C, Kraemer S, Lamason RL. 2025. *Rickettsia parkeri* forms extensive, stable contacts with the rough endoplasmic reticulum. J Cell Biol 224.

49. Goldstein JL, Brown MS. 1990. Regulation of the mevalonate pathway. Nature 343:425–30.

50. Budzinska A, Galganski L, Jarmuszkiewicz W. 2023. The bisphosphonates alendronate and zoledronate induce adaptations of aerobic metabolism in permanent human endothelial cells. Sci Rep 13:16205.

51. Ahyong V, Berdan CA, Burke TP, Nomura DK, Welch MD. 2019. A Metabolic Dependency for Host Isoprenoids in the Obligate Intracellular Pathogen *Rickettsia parkeri* Underlies a Sensitivity to the Statin Class of Host-Targeted Therapeutics. mSphere 4.

